# Cfap300 regulates the transdifferentiation of Corpuscle of Stannius cells in zebrafish

**DOI:** 10.1101/2025.10.06.680176

**Authors:** Usharani Nayak, Kalyani Sahoo, Praveen Barrodia, Rajeeb K. Swain

## Abstract

Stanniocalcin 1 (Stc1) is a hormone secreted by the Corpuscle of Stannius (CS) gland in teleost fish, including zebrafish, where it regulates calcium homeostasis. The CS gland forms by transdifferentiation of pronephric tubule epithelial cells at the distal early (DE) and distal late (DL) boundary. Although transdifferentiation is known in development and disease, the underlying mechanisms remain unclear. Here, we show that *cfap300*, a DNAAF implicated in primary ciliary dyskinesia (PCD), is essential for CS gland formation. TALEN-generated *cfap300* mutants develop normally and show no nephron segmentation defects, but exhibit impaired CS development, which could be partially rescued by *cfap300* mRNA injection. Further, *cfap300* mutants display increased expression of *cdh17* driven by Hnf1ba, which blocks CS precursor transdifferentiation. Knockdown of *cdh17* or *hnf1ba* restores CS formation. These results uncover a Cfap300–Hnf1ba–Cdh17 pathway that regulates epithelial-to-endocrine transdifferentiation, revealing a novel, lineage-specific role for a DNAAF protein in endocrine organogenesis.

## Introduction

Primary non-motile cilia are present on most mammalian cells and serve as sensory organelles, whereas the motile cilia are expressed in multiple organs where they function to move fluids and particles (1). Dynein arm assembly factors (DNAAF) are cytoplasmic proteins that are essential for the assembly and transport of dynein motor complexes, crucial for the movement of cilia (2). Mutations in DNAAF genes lead to primary ciliary dyskinesia (PCD), a group of rare genetic diseases with a diverse phenotype (3, 4). Zebrafish mutants of human DNAAF homologs have been shown to faithfully recapitulate the PCD phenotypes associated with human mutations, thus serving as genetic animal models of this group of diseases (5–7). Cilia and flagella-associated protein 300 (CFAP300, also known as DNAAF17) localises to the cytoplasm and cilia-associated compartment of motile ciliated cells (8). Biallelic loss-of-function mutations of CFAP300 have been linked to primary ciliary dyskinesia (PCD) in various populations, marked by situs inversus, respiratory cilia immotility, and loss of both the inner and outer dynein arms (IDA + ODA) (4, 8–10).

The organs containing motile cilia in zebrafish include, *inter alia,* the Kupffer’s vesicle, pronephros, olfactory organs and spinal central canal (11). Zebrafish pronephros consists of glomerulus, neck, proximal convoluted tubule (PCT), proximal straight tubule (PST), distal early (DE), distal late (DL) and pronephric duct (PD). The proximal segments are homologous to the PCT and PST in mammals, while the distal segments are homologous to the mammalian thick ascending limb (TAL) and distal convoluted tubule (DCT), respectively (12). The zebrafish pronephros is lined by both mono and multi-ciliated cells, which beat in a coordinated manner to generate fluid flow through the pronephric ducts. This ciliary-driven flow is essential for waste excretion, the prevention of fluid build-up. A pronephric cilia defect often leads to a pronephric cyst in zebrafish (5, 6). The zebrafish pronephros also possesses the site of origin of an endocrine gland called the Corpuscle of Stannius (CS), which is unique to the teleost class and maintains calcium homeostasis through the secretion of stc1 hormone (13). CS gland-forming cells originate by transdifferentiation of the pronephric epithelium present between DE and DL segments and gradually extrude out to form a paired, independent gland (14). Transdifferentiation refers to the process by which a fully differentiated, mature somatic cell transforms into a different cell type. This process can occur through a direct conversion from one cell type to another or might involve a stage of dedifferentiation before the final cell fate is achieved (15). In zebrafish, several transcription factors (e.g. Hnf1b, Irx2a/3b, Tbx2a/b, Sim1a) and signalling pathways (e.g. Notch, RA and Fgf) are known to control CS gland development (12, 14, 16–19). Their specific functions during transdifferentiation and extrusion, however, remain to be elucidated.

In this study, we report that zebrafish *cfap300* is expressed in the pronephros and multiple other organs that possess motile cilia. We have studied the function of *cfap300* by creating a mutant zebrafish using TALEN. These mutants show no obvious morphological abnormalities, grow like wild type (WT) and are fertile. They show normal nephron segmentation compared to WT, but the CS gland development is impaired in these mutants. We show that impaired CS gland formation is due to the lack of transdifferentiation of distal pronephric epithelia to CS gland-forming cells. *cfap300* mRNA injection in *cfap300* mutants restored CS development, confirming its role in CS gland development. As cell adhesion plays an important role in differentiation, the level of cadherins expressed in the pronephros was checked in WT and *cfap300* mutants, and it was found that *cdh17* is overexpressed in these mutants compared to WT. We further show that *hnf1ba* is significantly up-regulated in these mutants and could drive the expression of *cdh17*. Knockdown of *hnf1ba* or *cdh17* rescued the transdifferentiation and CS gland formation defect in *cfap300* mutants. These data suggest that *cfap300*, a gene implicated in PCD, regulates transdifferentiation of CS cells from pronephric epithelium through *hnf1ba* and *cdh17*, thus unveiling a novel role for a DNAAF family member.

## Results

### *cfap300* is expressed in the pronephros and motile cilia-bearing tissues during zebrafish embryogenesis

The CFAP300 is highly conserved among vertebrates and its orthologs are present in most species containing motile cilia, including invertebrates (8, 20). Whole-mount in situ hybridization (WISH) was carried out to determine the expression of *cfap300* mRNA during zebrafish embryogenesis. The expression of *cfap300* was not detected from the 2-cell to the 6 hpf stage (Figure 1a-d). Expression of *cfap300* was observed in the Kupffer’s vesicle at 10 hpf, a transient organ containing ciliated cells that is essential for determining left-right asymmetry (Figure 1e). *cfap300* is expressed in the notochord (NC) and intermediate mesoderm (IM) at 14 hpf (Figure 1f). Its expression in pronephros (P), olfactory placodes (OP), notochord and tail bud (TB) was detected at 22 hpf (Figure 1g). In 24 hpf embryos, *cfap300* is expressed in the pronephros, olfactory placodes, notochord, epiphysis (E), tegmentum (T), floor plate (FP) and tail bud (Figure 1h). *cfap300* is not expressed in the glomerulus, but its expression starts at the neck segments of the pronephros. Strong expression of *cfap300* can be seen in the whole pronephric tubule and duct (Figure 1i). Histological analysis confirmed its expression in the pronephros (Figure 1j). The expression of *cfap300* in the above tissues remained up to 48 hpf (Figure 1k-m). Two-colour WISH of *cfap300* along with *pdzk1* confirmed its expression in the proximal and distal early parts of the tubule and *slc12a3* confirmed its expression in the distal late segment (Figure 1n-o). The expression pattern of *cfap300* is in agreement with its proposed role in cilia biogenesis and function because organs like Kupffer’s vesicle, pronephros, olfactory placode, tegmentum and epiphysis contain motile cilia (11, 21). Its expression in the entire pronephros tubule and duct, and not just the motile cilia-containing region, suggests that it may regulate tissue-specific developmental programs in the kidney in addition to its proposed role in cilia.

**Figure 1:**
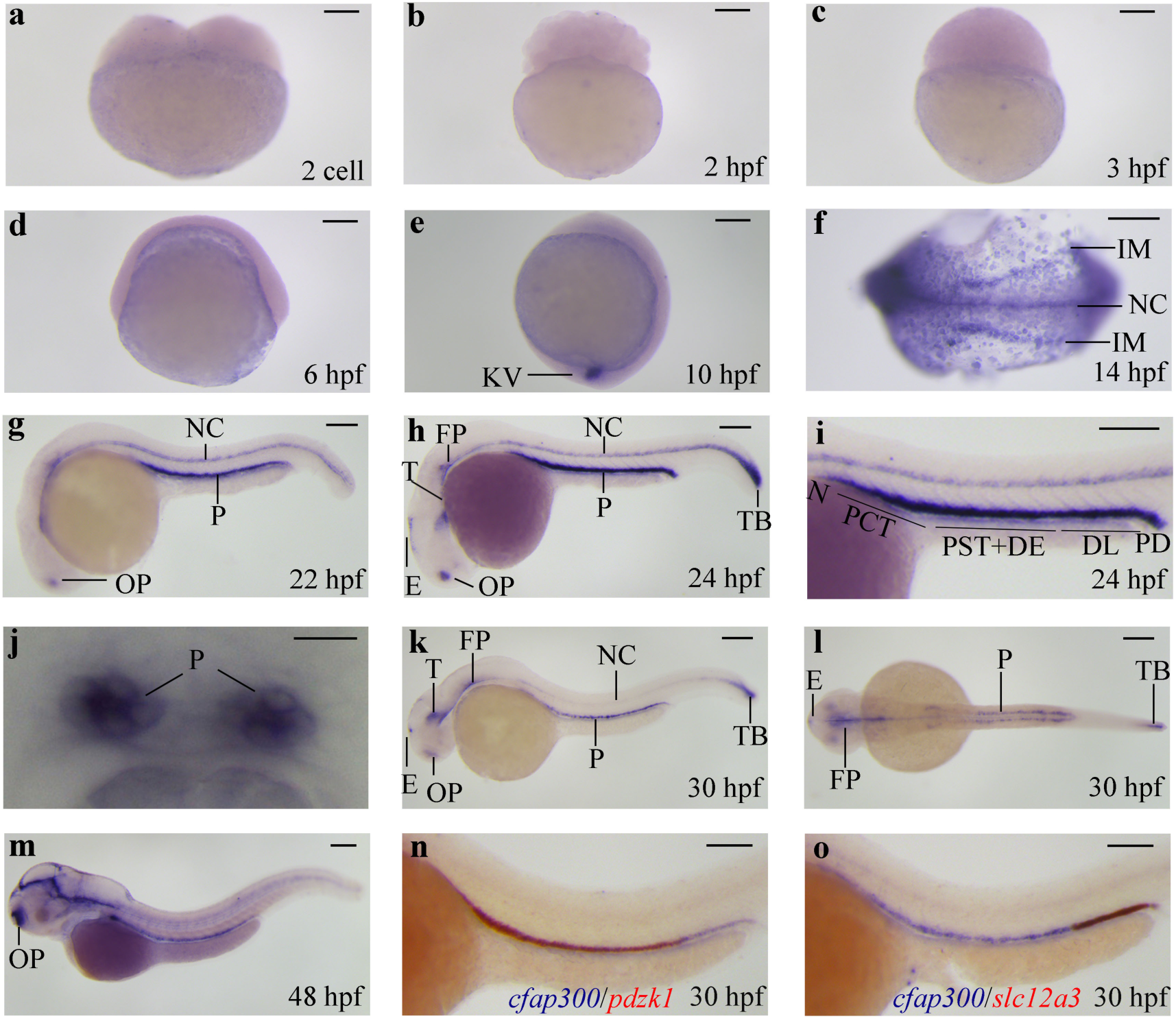
*cfap300* is expressed in the pronephros and other motile cilia-bearing tissues during zebrafish development. WISH data show that *cfap300* is not expressed until 10 hpf **(a-d)**. *cfap300* is expressed in Kupffer’s vesicle (KV) at 10 hpf **(e)** and in notochord (NC) and intermediate mesoderm (IM) at 14 hpf (**f)**. It is expressed in the pronephros (P) and the olfactory placodes (OP) at 22 hpf **(g)**; in epiphysis (E), tegmentum (T), floor plate (FP) and tail bud (TB) in addition to NC and OP at 24 hpf **(h)**. Close observation shows that *cfap300* is expressed in all segments of the pronephros tubule and duct except the glomerulus **(i)**, and histology confirms its expression in pronephros **(j)**. *cfap300* expression persists in the above tissues at least until 48 hpf **(k-m)**. Two colour WISH of *pdzk1* confirmed *cfap300* expression in the pronephric tubule and *slc12a3* confirmed its expression in the duct **(n and o)**. hpf: hours post fertilisation. Scale bar 150 μm, except 1j in which it is 25 μm.

### Corpuscle of Stannius gland formation is impaired in *cfap300* mutants and morphants

A TALEN-mediated *cfap300* mutant zebrafish was generated by targeting exon 2. Zebrafish mutant fish with deletion of 26 nucleotides in the coding region (Δ26) were confirmed by Sanger sequencing and homozygous mutants (henceforth referred to as *cfap300^−/−^*or mutants) were used in all experiments (Figure 2A). High-resolution melt curve (HRM) and heteroduplex analysis (HD) were used to identify the mutant allele of *cfap300* (Figure 2B and C). Deletion of 26-nucleotides in *cfap300* introduces a premature termination codon (PTC) after 55 amino acids of Cfap300 and a leucine in place of a phenylalanine. It is well known that PTC can lead to nonsense-mediated decay of the mRNA (22). Hence, we checked if the *cfap300* mRNA undergoes NMD and is degraded in *cfap300^−/−^* embryos. RT-qPCR and WISH were carried out on wild type (WT) and *cfap300^−/−^* embryos, which showed that the *cfap300* mRNA is degraded in the mutants, thus indicating that the mutants will have no functional Cfap300 protein (Figure 2D and E).

**Figure 2:**
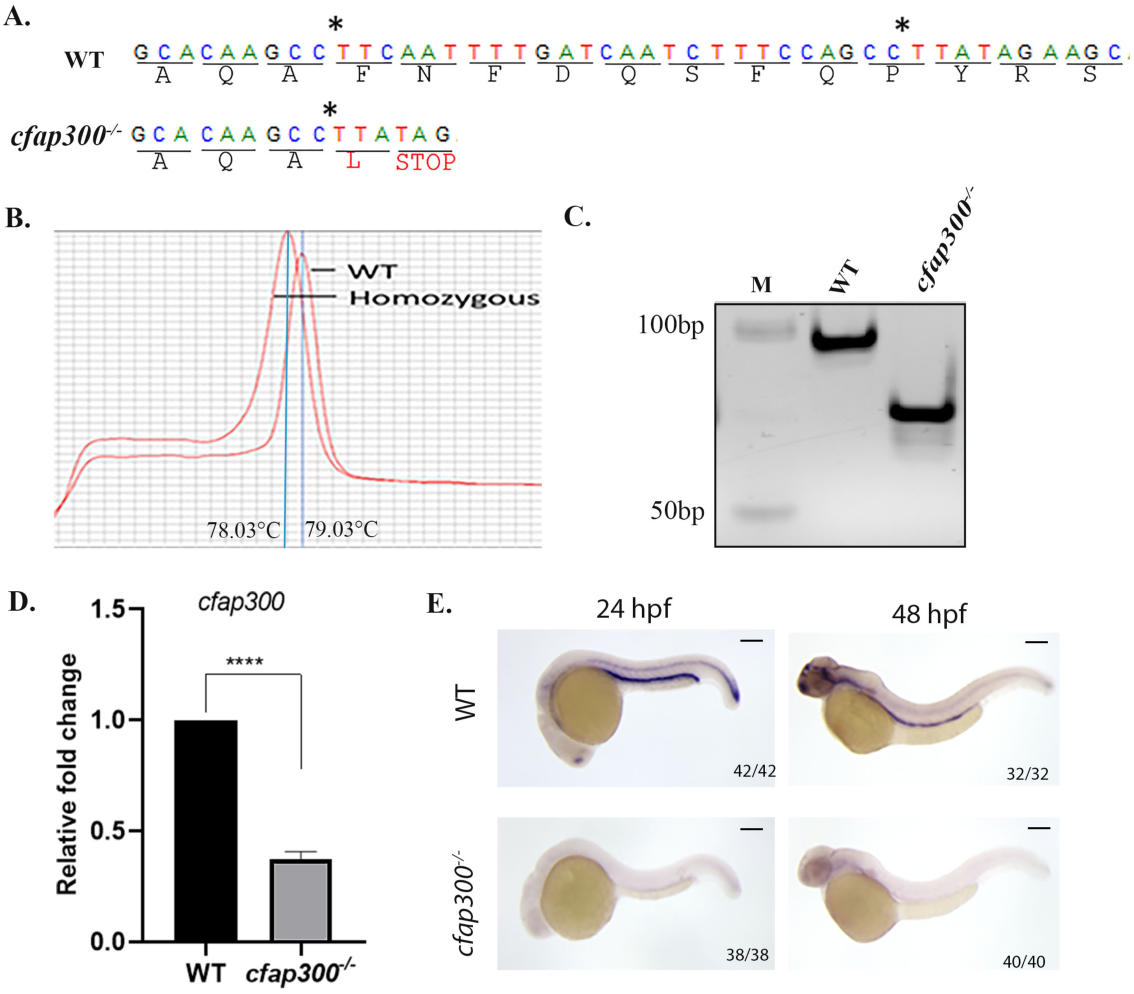
Generation of the *cfap300* loss-of-function mutant. **(A)** Sanger sequencing reveals a 26-nucleotide deletion in the *Cfap300* coding region, resulting in a premature termination codon (PTC) after 55 amino acids of Cfap300 and a Leucine. **(B and C)** High-resolution melt curve and heteroduplex analysis to identify the mutant allele of *cfap300*. (**D**) RT-qPCR analysis of RNA extracted from pooled 24 hpf WT or *cfap300^−/−^* embryo shows NMD-mediated degradation of *cfap300*. p<0.0001 from unpaired t-test. (**E)** WISH shows decreased *cfap300* transcripts in *cfap300^−/−^* embryos, confirming that *cfap300* mRNA is unstable. Scale bar 150 μm.

The *cfap300* mutants do not exhibit gross morphological abnormalities at 24 hpf or later (data not shown). Since *cfap300* is strongly expressed in the pronephros, we checked if pronephros development is impaired in the *cfap300^−/−^* embryos. Zebrafish pronephros segments can be distinguished by analysing the expression of marker genes that are expressed in a segment-specific manner (12). The somite marker *xirp2a* was used to measure the pronephros segment length with respect to the somite number. In wild-type embryos, *sodium/inorganic phosphate symporter* (*slc20a1a*) is expressed in part of the nephron adjacent to the 3^rd^ to 7^th^ somite in 48 hpf embryos, which demarcates the proximal convoluted tubule (PCT). *Transient receptor potential cation channel*, *subfamily M, member 7* (*trpm7*) is expressed in the proximal straight tubule (PST) aligned with the 8^th^ to 11^th^ somite. *Sodium/potassium/chloride transporter slc12a1* is expressed in part of the nephron adjacent to the 12^th^ and 14^th^ somite at 48 hpf, demarcating the distal early (DE) segment of the nephron. *Sodium/chloride transporter slc12a3* is expressed in part of the nephron adjacent to the 15^th^ and 17^th^ somite, demarcating the distal late (DL) segment of the nephron (23). The expression levels and domains of *slc20a1a*, *trpm7*, *slc12a1* and *slc12a3* were identical in both WT and *cfap300*^−/−^ embryos at 48 hpf (Figure 3A), indicating that *cfap300* may not regulate nephron segmentation. The expression domain of *stc1* that demarcates the Corpuscle of Stannius was not changed. However, the *stc1* mRNA level was greatly reduced in the *cfap300* mutants, indicating that *cfap300* may have a role in CS development or *stc1* expression (Figure 3A). We also examined multiciliated cell (MCC) development in the pronephros by analysing the ciliogenesis markers *odf3* and *rfx2*. Their expression domains and intensity were indistinguishable between wild-type and mutant embryos (Figure 3B and C). Thus, *cfap300* mutants do not display broad defects in MCC formation, reinforcing the idea that its role is specific to CS gland development.

**Figure 3:**
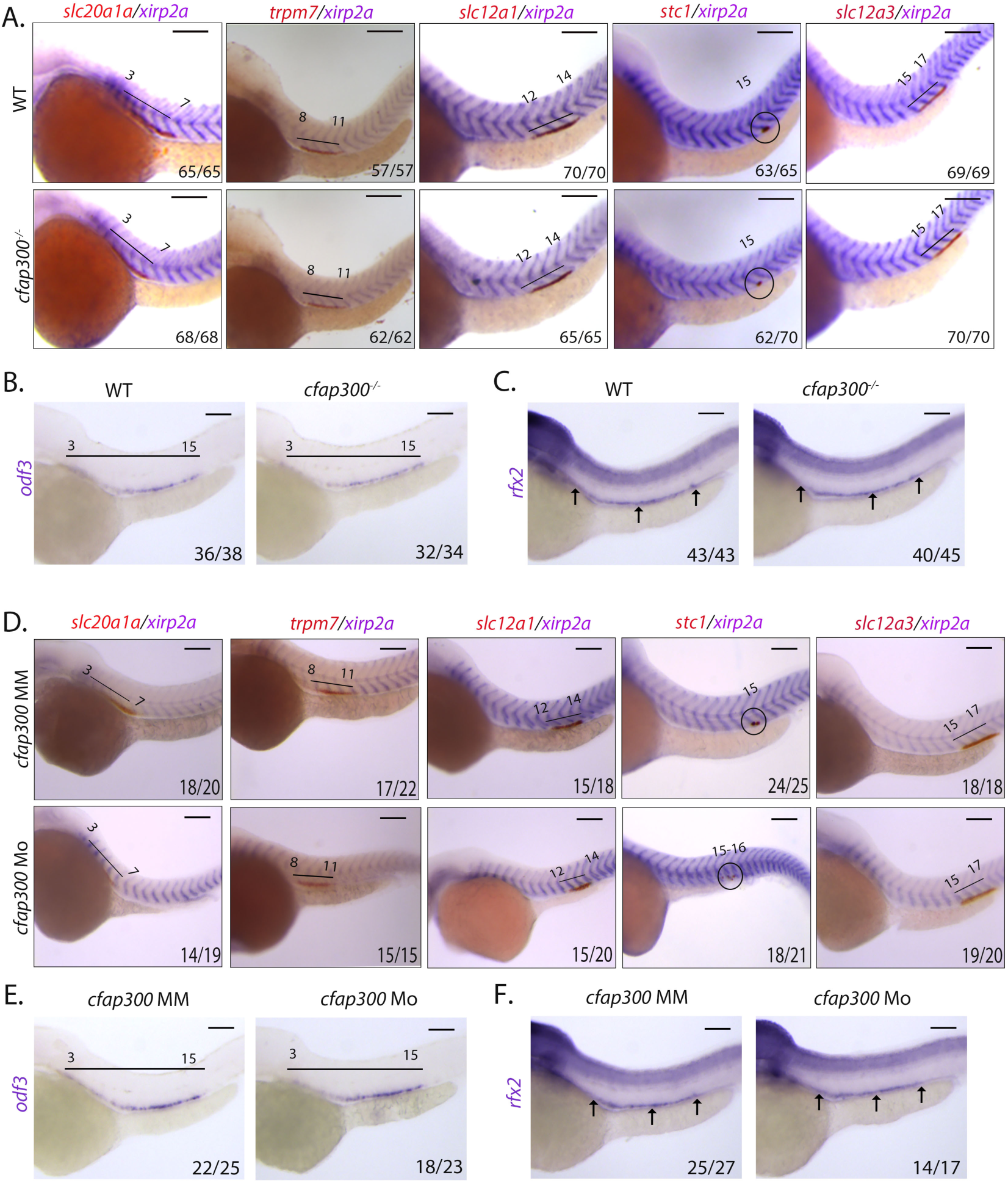
Pronephros segmentation and MCC formation are not affected in *cfap300* mutants or morphants. (**A**) WISH with pronephros segment-specific markers *slc20a1a* (PCT), or *trpm7* (PST), or *slc12a1* (DE), or *slc12a3* (DL) or *stc1* (CS) with a somite marker *xirp2a* shows reduced *stc1* expression, without any change in pronephros segmentation in *cfap300* mutants. (**B and C)** WISH for *odf3* or *rfx2* shows normal MCC formation in *cfap300* mutants. (**D)** WISH against pronephros segment-specific markers with a somite marker on the control mismatch (*cfap300*-MM) and translation blocking (*cfap300*-MO) antisense morpholino-injected embryos shows reduced *stc1* expression without alteration in pronephros segmentation in *cfap300* morphants. **(E and F)** Unaltered MCC formation in *cfap300* morphants. Scale bar 150 μm.

It has been reported that nonsense-mediated decay of mRNA can induce transcriptional compensation, whereas antisense morpholino oligo-mediated knockdown has no such effect (24). We hypothesised that the lack of any major defect in nephron segmentation and MCC formation may be a result of the upregulation of other genes in the *cfap300* mutants that can compensate for its loss of function. Hence, we checked if morpholino-mediated knockdown of *cfap300* results in defects not seen in the *cfap300^−/−^*embryos. A translation blocking *cfap300* morpholino (*cfap300*-MO) and a negative control mismatch morpholino (*cfap300*-MM) were injected into 1-cell-stage zebrafish embryos and the effect of these morpholinos on morphology, nephron segmentation and MCC formation was analysed. The gross morphology or nephron segmentation or MCC formation was normal in the morphants (Figure 3D-F). The *stc1* expression, however, was decreased in the morphants, recapitulating the phenotype seen in the mutants (Figure 3D). These data indicate that *cfap300* specifically regulates CS development without broadly impacting nephron segmentation or ciliogenesis.

To confirm that *cfap300* regulates CS gland formation and not just *stc1* expression, we analysed the CS gland development using multiple markers that recognise this gland (12, 25). *fgf23* and *gata3* expression were greatly reduced in the *cfap300^−/−^*embryos, indicating that indeed CS gland formation is impaired in the mutants (Figure 4A). Interestingly, *gata3* expression in other anatomical structures, such as spinal cord neurons and pronephric duct was unaffected, further strengthening the idea that the *cfap300* effect is limited to CS gland formation (Figure 4A).

**Figure 4:**
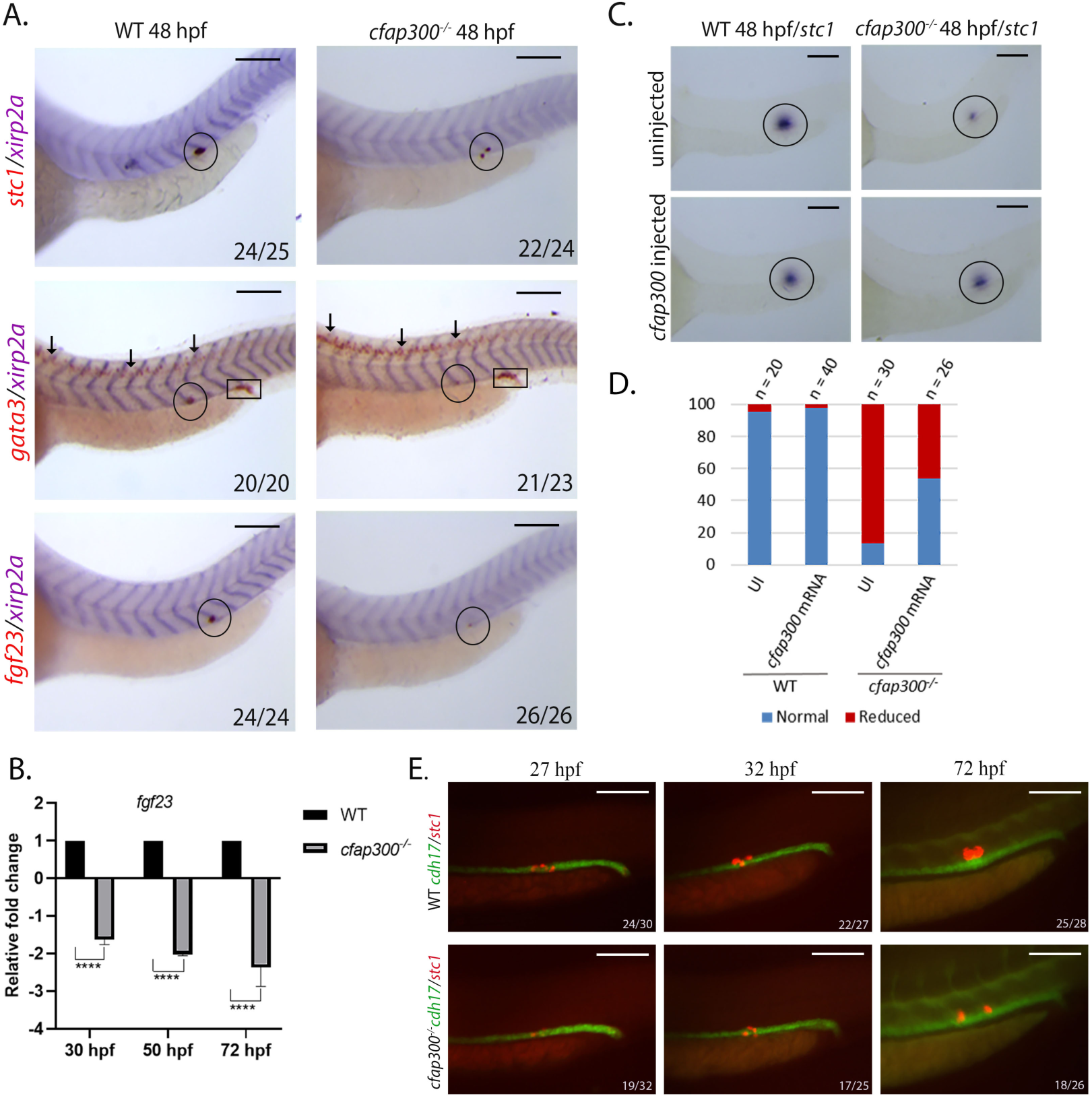
CS gland formation is impaired in *cfap300* mutants. **(A)** WISH showed decreased CS gland marker gene expression of *stc1*, *gata3* and *fgf23* (black circles) in *cfap300* mutants. Notably, *gata3* expression outside the CS gland, including spinal cord (black arrows) and pronephric duct (black box), remains unaffected. **(B)** RT-digital PCR analysis identified decreased *fgf23* expression (p<0.0001, 2-way ANOVA analysis) during the CS gland morphogenesis (30, 50, and 72 hpf). **(C)** *cfap300* mRNA injection restores *stc1* expression in *cfap300* mutants. **(D)** Quantification showed *cfap300* mRNA injection resulting in normal *stc1 expression in* ∼55% *cfap300^−/−^* embryos, while only ∼10% of control-uninjected (UI) *cfap300^−/−^* embryos show normal *stc1* expression (p<0.001, Cochran–Mantel–Haenszel (CMH) test). **(E)** Fluorescent WISH showed reduced *stc1*-positive CS cells in *cfap300* mutants during 27-72 hpf. The *cdh17* expression (green) marks the entire pronephric tubule and duct, whereas *stc1* (red) marks the CS gland. Scale bar 150 μm.

*fgf23* expression was analysed by digital PCR on cDNA prepared from different stages of CS gland formation to confirm the WISH data. We found that *fgf23* mRNA level was greatly reduced in the *cfap300* mutants compared to the WT embryos during different stages of CS gland formation (Figure 4B). Moreover, microinjection of *cfap300* mRNA into the *cfap300* mutants resulted in 55% embryos having normal *stc1* expression comparable to WT embryos. These data establish that loss of *cfap300* leads to impaired CS gland development that can be rescued by expression of *cfap300* mRNA (Figure 4C and D; Supplementary Figure 1). These findings provide the first evidence that a conserved ciliary gene such as *cfap300*, previously associated only with motile cilia function and ciliopathies, has a lineage-specific role in regulating endocrine organ formation.

### *cfap300* is necessary for CS gland cell transdifferentiation

A defined number of cells from the pronephros tubule bordering the DE and DL segments transdifferentiate (completed by 30 hpf) to form the CS cells, undergo apical constriction and extrude from the tubule and form the CS gland by 72 hpf. To check how *cfap300* might take part in this gland formation, a two-colour fluorescent WISH was carried out using *cdh17* as a marker for the pronephric tubule and *stc1* as a marker for CS cells. The number of CS cells at 27 hpf in *cfap300* mutants was greatly reduced compared to the wild-type embryos (Figure 4E). The fluorescent WISH carried out at 32 and 72 hpf indicated that the transdifferentiation is impaired and not delayed in the mutants (Figure 4E). We also observed that a low number of CS cells that form in the *cfap300^−/−^*embryos exhibit incomplete extrusion out of the nephron epithelium (Figure 4E). These results demonstrate that *cfap300* is required for epithelial to endocrine transdifferentiation underlying CS gland formation, a previously unrecognized mechanism in vertebrate kidney and endocrine development.

### The impaired CS gland formation in *cfap300* mutants is due to upregulation of *cdh17*

It is reported that *cdh17*, which is expressed in the tubule and duct of zebrafish pronephros, is downregulated in the border between the DE and DL (14). We reasoned that *cdh17* and other cadherins may be involved in the process of transdifferentiation. Hence, we checked the status of cadherins expressed in the pronephros of zebrafish (*cdh1*, *cdh6* and *cdh17*) by RT-qPCR at 48 hpf. All cadherins except *cdh17* showed non-significant change of expression, whereas *cdh17* expression was significantly upregulated in *cfap300^−/−^* embryos (Figure 5A). Then the *cdh17* mRNA levels in the pronephros during different stages of CS gland development, such as end of transdifferentiation (30 hpf), extrusion (50 hpf) and expansion of CS cells (72 hpf) were checked using RT – digital PCR and WISH. The *cdh17* was significantly upregulated in all three stages and the WISH results showed the overexpression is throughout the tubule (Figure 5B and C). This indicates that the upregulation of *cdh17* in the pronephros of zebrafish may be the reason behind impaired transdifferentiation of CS cells from the tubule of the pronephros, leading to CS gland formation defect in *cfap300* mutants. If *cdh17* upregulation leads to impaired CS gland formation, we reasoned that the knockdown of *cdh17* in *cfap300* mutants would restore the CS gland formation. We took advantage of the F0 crispant method to test this hypothesis. Two guide RNAs (gRNA) were designed targeting the exon-4 and exon-8 of zebrafish *cdh17* and were tested for their efficacy (Supplementary Figure 2). The microinjection of *cdh17* gRNA led to the NMD-mediated degradation of *cdh17* mRNA in these crispants, suggesting that the *cdh17* crispants will lack Cdh17 protein (Figure 6A). We then checked CS gland formation in *cdh17* crispants at 30, 50 and 72 hpf by analyzing *stc1* expression. The CS gland formation was largely restored in the *cdh17* crispants created in the *cfap300^−/−^* background (Figure 6B). Only 10% of *cfap300^−/−^*embryos had normal *stc1* expression at 30 hpf, whereas 60% embryos showed normal CS development in *cdh17* crispants created in these mutants (Figure 6C). Similar results were obtained when *cdh17* crispants created in the *cfap300^−/−^*background were analyzed at 50 and 72 hpf (Figure 6C). These experimental observations support the hypothesis that upregulation of *cdh17* in *cfap300* mutants leads to impaired transdifferentiation and reduced CS gland formation, which can be restored by downregulating *cdh17*. These data identify a mechanism whereby persistent *cdh17* expression acts as a barrier to CS cell transdifferentiation, linking *cfap300* function to an adhesion molecule regulation.

**Figure 5:**
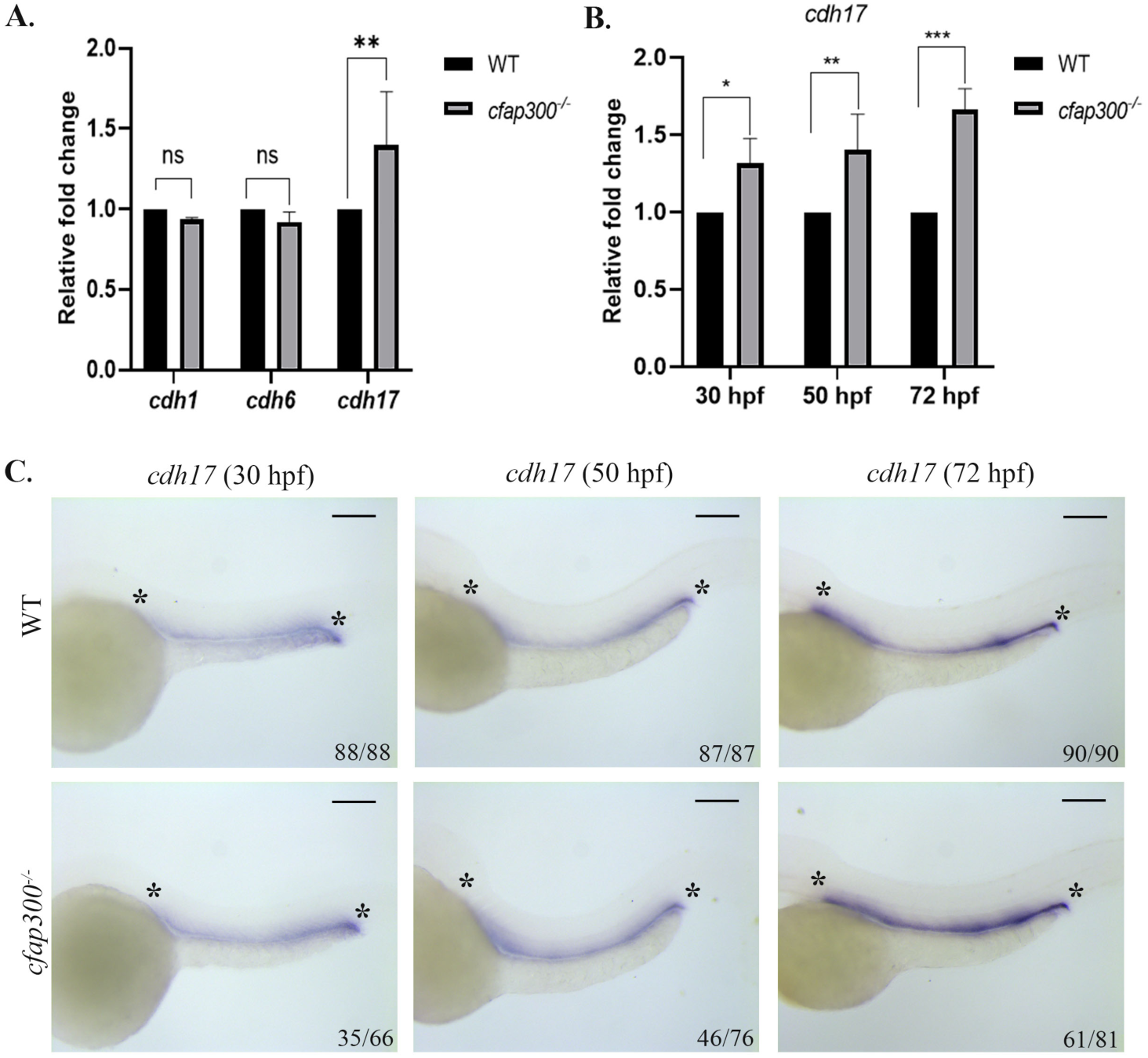
*cdh17* expression is upregulated in *cfap300* mutants. (**A**) RT-qPCR analysis of *cdh1*, *cdh6* and *cdh17* at 48 hpf. Only *cdh17* expression was significantly increased in *cfap300* mutants (\*\***/**p=0.0057, 2-way ANOVA). **(B)** RT – digital PCR confirms increased *cdh17* expression at different stages of CS development in *cfap300* mutants (*/p<0.03, **/p<0.007 and ***/p=0.0001, 2-way ANOVA). **(C)** WISH showed increased *cdh17* expression in ∼53%, 60% and 75% embryos at 30, 50, and 72 hpf *cfap300^−/−^* embryos, respectively. Scale bar 150 μm.

**Figure 6:**
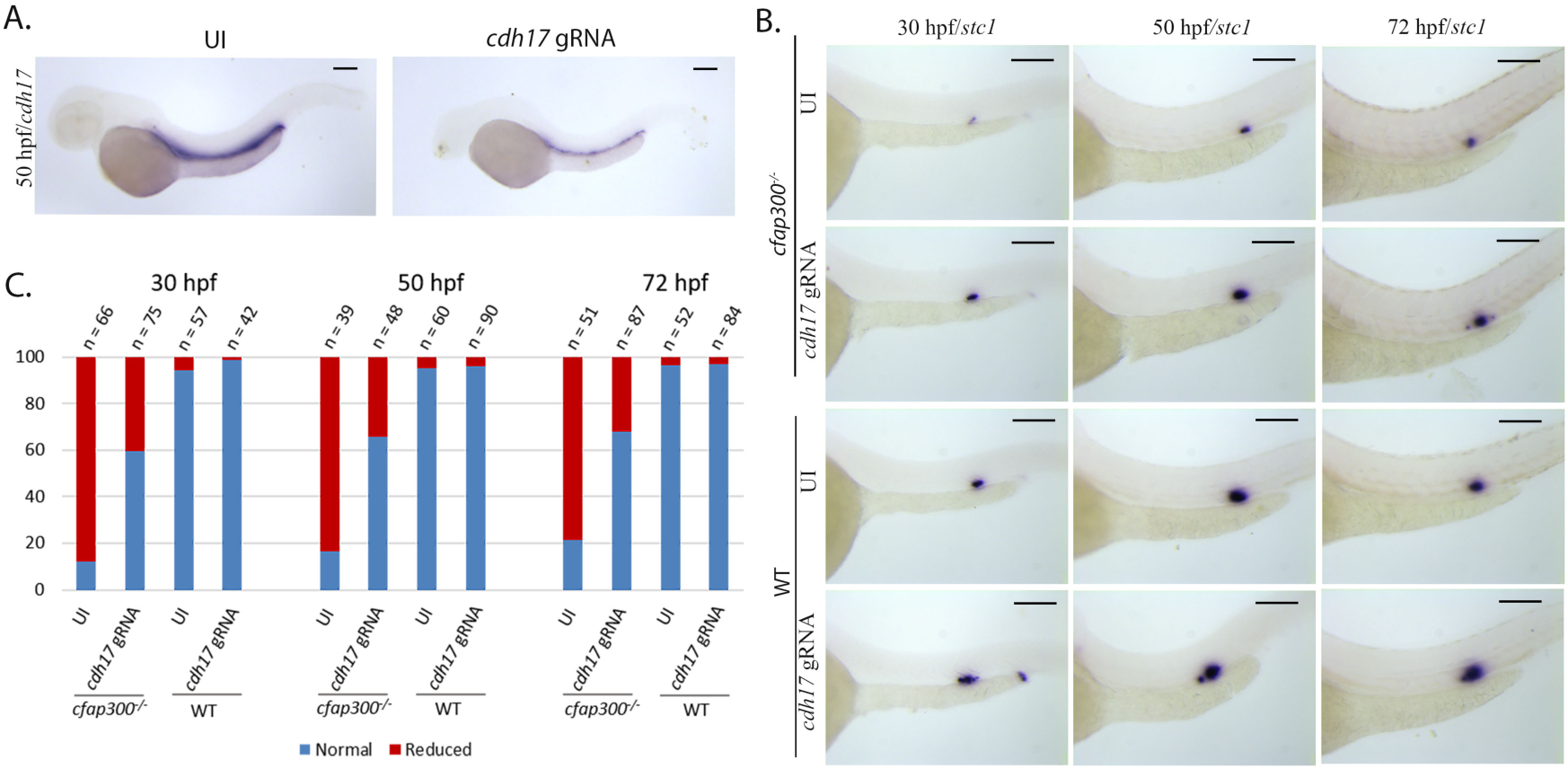
*cdh17* knockdown restores CS gland formation in *cfap300* mutants. (**A**) WISH showed decreased *cdh17* expression in *cdh17* crispants, indicating NMD-mediated *cdh17* mRNA degradation. **(B)** WISH against *stc1* showed restored CS gland development in *cdh17* crispants in *cfap300* mutants. **(C)** Quantification showed ∼60% *cfap300^−/−^/cdh17* crispants embryos had normal *stc1* expression compared to ∼10% in *cfap300*^−/−^ at 30, 50 and 72 hpf. The statistical significance of recovery in the mutant was determined using the Cochran–Mantel– Haenszel (CMH) test (p<0.001). Scale bar 150 μm.

### *hnf1b*a transcription factor is upregulated in the *cfap300* mutants and regulates *cdh17* expression

We next investigated how *cdh17* becomes misregulated in *cfap300* mutants. The 5kb upstream sequence of *cdh17* that is reported to drive its expression in the zebrafish pronephros was analysed for transcription factors that can bind to this region using Alggen software (http://www.lsi.upc.es/~alggen) (26, 27). Two hundred and thirty-four transcription factors were identified that can potentially bind to this region. Spatial expression of these TFs during zebrafish development was checked at the Zfin.org database to identify TFs that are expressed in a spatially overlapping manner with the *cdh17* in the zebrafish pronephros. Zebrafish *cjun*, *ppargc1a*, *hnf1ba*, *hnf1bb* and *c-myc* were found to be expressed in a complete or partially overlapped fashion with *cdh17*. The expression of these TF was analyzed in the *cfap300* mutant embryos at 48 hpf by RT-qPCR. *c-myc* expression was downregulated and *hnfb1a* expression was upregulated in the *cfap300* mutants, whereas the expression of other genes was not altered (Figure 7A). Next, we checked the ability of *c-myc* and *hnf1ba* to regulate *cdh17* transcription by creating crispants (*c-myc*) or morphants (*hnf1ba*). Since *hnf1bb* is a paralog of *hnf1ba*, it was also taken, although its expression was not altered in the mutants. Our analysis showed that *hnf1bb* crispants had normal, *c-myc* crispants showed enhanced and *hnf1ba* morphants showed reduced *cdh17* expression compared to WT embryos at 48 hpf (Figure 7B). Since *hnf1ba* expression is upregulated in *cfap300* mutants and could enhance *cdh17* transcription, we checked its level of expression at different stages of CS development. The RT – digital PCR and WISH on WT and *cfap300* mutant embryos between 30 and 72 hpf show that *hnf1ba* transcription is upregulated in the mutant embryos during the CS gland formation (Figure 7C and D). These data support the hypothesis that *hnf1ba* and its transcriptional target *cdh17* are upregulated in the *cfap300* mutants, which results in impaired CS gland formation.

**Figure 7:**
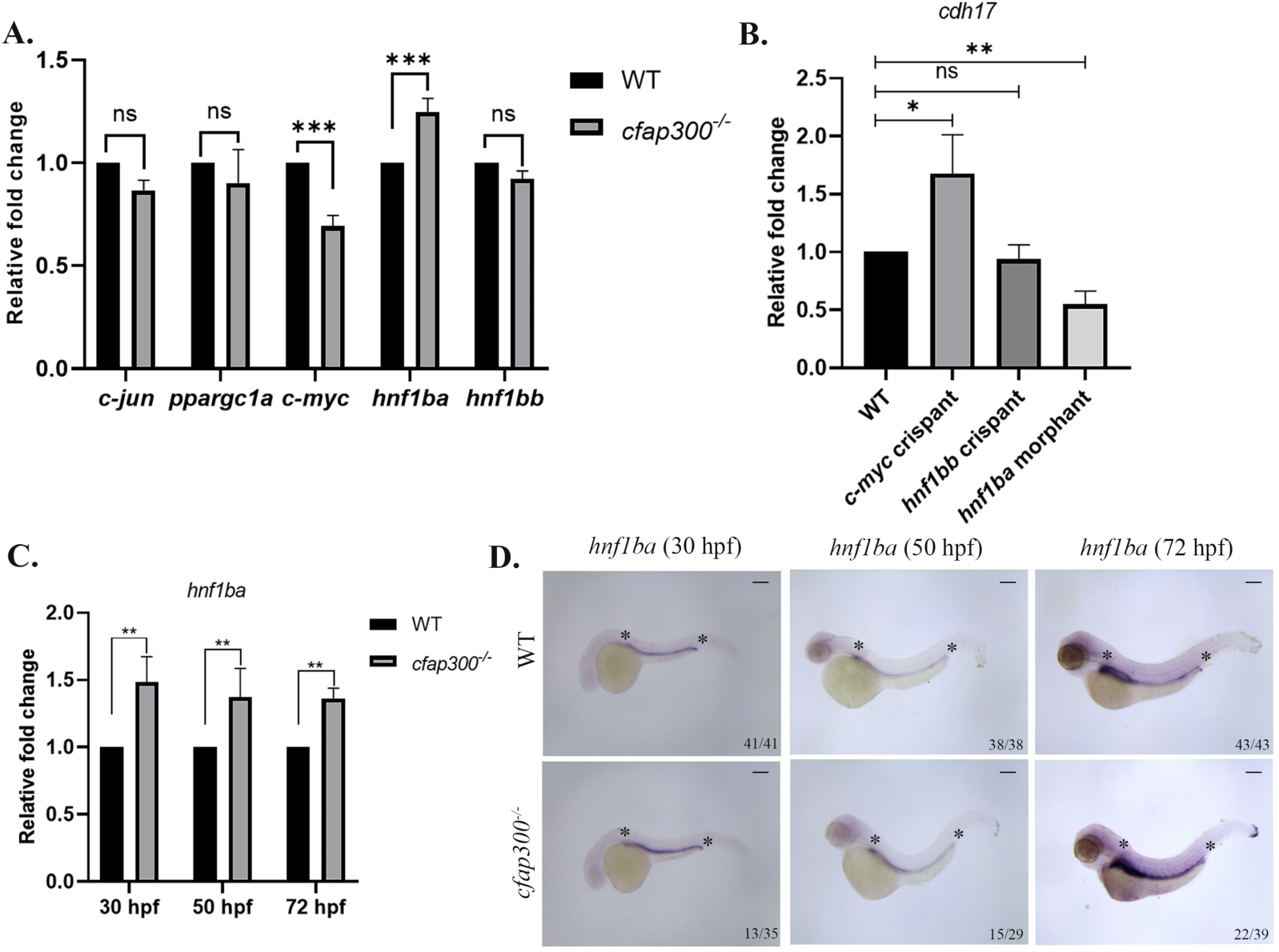
Cfap300 negatively regulates *hnf1ba* but not *hnf1bb* expression in developing pronephros. **(A)** Quantification of the transcripts of transcription factors that are predicted to bind to the regulatory sequence of *cdh17* and are known to be expressed in the pronephros. Decreased *c-myc* and increased *hnf1ba* transcripts in *cfap300* mutants compared to wild-type embryos (***/p<.001, 2-way ANOVA). **(B)** Quantification of *cdh17* transcripts in c-*myc* or *hnf1bb* crispants or *hnf1ba* morphants at 48 hpf (*/p=0.02, **/p=0.002, unpaired t-test). **(C)** Comparative analysis of *hnf1ba* transcripts in embryos during different stages of CS gland formation (30 – 72 hpf). (D) Lateral views of WISH embryos against *hnf1ba*. *hnf1ba* is increased in *cfap300* mutants during CS gland development. Statistical significance determined by Student’s t-test (**) p<0.01. Scale bar 150 μm.

### Knockdown of *hnf1ba* partially restores CS development in *cfap300* mutants

We have provided evidence that *hnf1ba* is upregulated in the pronephros of *cfap300* mutants, leading to upregulation of *cdh17*, impaired transdifferentiation of CS cells and defective CS gland formation. If this is true, then downregulation of *hnf1ba* in *cfap300^−/−^* embryos should lead to restoration of CS gland formation. To test this pathway directly, we knocked down *hnf1ba* in *cfap300^−/−^* embryos using a translation-blocking antisense morpholino. Hnf1ba downregulation led to reduced *cdh17* expression and partial restoration of CS formation, with over 80% of *cfap300* mutant embryos at 30, 50, and 72 hpf showing normal or elevated *stc1* expression (Figure 8A and B; Supplementary Figure 3). This functional rescue confirms that *hnf1ba* acts upstream of *cdh17* and positions *cfap300* as a key regulator of CS gland development.

**Figure 8:**
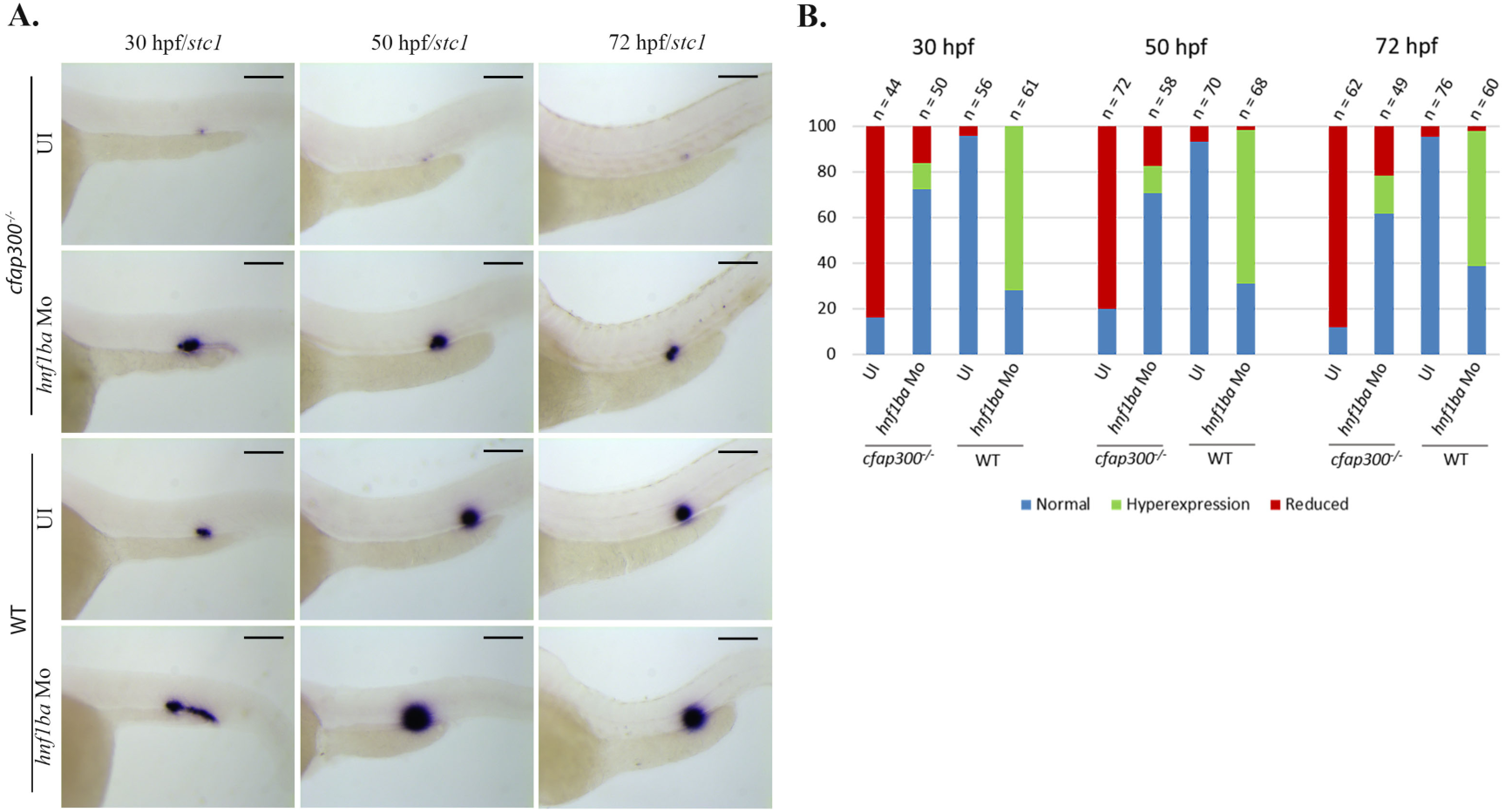
Knockdown of *hnf1ba* restores the CS development in *cfap300* mutants. **(A)** Lateral views of whole mount *in situ* hybridised wild-type or *cfap300*^−/−^ embryos, either control or a translation-blocking morpholino against *hnfba* injected. *hnf1ba* knockdown largely restored the *stc1*-expressing CS gland cells in the *cfap300* mutants. (**B)** Bar graph quantifies the percentage of embryos with *stc1* expression categorised as normal, ectopic or reduced in uninjected (UI) or *hnf1ba* morpholino-injected *cfap300* mutant or WT. *hnf1ba* knockdown in *cfap300* mutants restored *stc1* expressing CS cells in ∼50-60% embryos. The statistical significance of recovery in the mutant was determined using the Cochran-Mantel-Haenszel test (p<0.001). Scale bar 150 μm.

## Discussion

The Dynein axonemal assembly factors (DNAAFs) are well-known essential proteins involved in the assembly of axonemal dynein arms and cilia biogenesis (4). Apart from these canonical functions, DNAAF also contributes to cell division and cell fate determination (28–31). In this study, we report that Cfap300, a protein classified as a DNAAF, regulates CS gland morphogenesis by suppressing *hnf1ba* and its target *cdh17* expression in zebrafish. Several outcomes of this study support the claim. *cfap300* is predominantly expressed in the pronephric tubule during CS gland morphogenesis and *cfap300* mutants showed reduced CS glands. These mutants show increased *hnf1ba* and *cdh17* expression and the CS gland is restored by downregulation of *hnf1ba* or *cdh17* in the mutants. Cadherins, including *cdh1*, *cdh6, cdh16,* and *cdh17,* are known to be expressed in the developing zebrafish pronephros. Interestingly, our study found pronephric expression of *cdh17* is regulated by Cfap300, while *cdh1*, *cdh6* expression remain unaltered in *cfap300* mutants. The function of CDH17 in cell adhesion and cell signalling, including the Wnt/β-catenin, NF-κB, α2β1 integrin, FAK, and Ras pathways, has been well documented (32–36). Since our study identified reduced CS gland along with increased *cdh17* expression in *cfap300* mutants, we speculate that downregulation of Cdh17 is necessary for the transdifferentiation of the pronephric epithelial cells to CS cells. A recent report showed that *cdh16* is expressed in most segments of the pronephros but primarily localises to the CS gland at 48 hpf. The *cdh16* mutants show impaired acoustic sensory gating in zebrafish as a result of a defect in calcium homeostasis (37). Thus, it will be interesting to explore the role of Cdh16 in Cfap300-mediated CS development in future.

Hnf1ba and Hnf1bb are two zebrafish paralogues of human HNF1B, expressed in many organs, including pronephros, liver, pancreas and gut (38–40). In the pronephros, *hnf1ba* is expressed broadly, whereas *hnf1bb* expression is restricted to proximal and distal early segments (38). Despite the differences in their expression domains, they have been proposed to carry out similar functions in zebrafish pronephros segmentation. We show here that *hnf1ba* and not *hnf1bb* is upregulated in the *cfap300* mutants, and suppression of *hnf1ba* in *cfap300* mutants resulted in normal CS development without alteration in pronephros morphogenesis, indicating that Hnf1ba is essential for CS gland development, and dispensable in nephron segmentation. Conversely, Hnf1bb, which has been shown to regulate nephron segmentation, may not regulate CS development (26, 27). It has been reported that Hnf1b is translocated from the nucleus to the cytoplasm in the cells between the DE and DL boundary cells of the zebrafish (14). As this study shows that suppression of *hnf1ba* or *cdh17* in *cfap300* mutants rescued CS gland morphogenesis phenotype, and *cdh17* is a transcriptional target of Hnf1ba, one could reason that the downregulation of Hnf1ba target genes is necessary for the transdifferentiation of CS forming cells from the nephric epithelium. In the future, it will be important to know whether other targets of Hnf1ba are also involved in this process (Figure 9).

**Figure 9:**
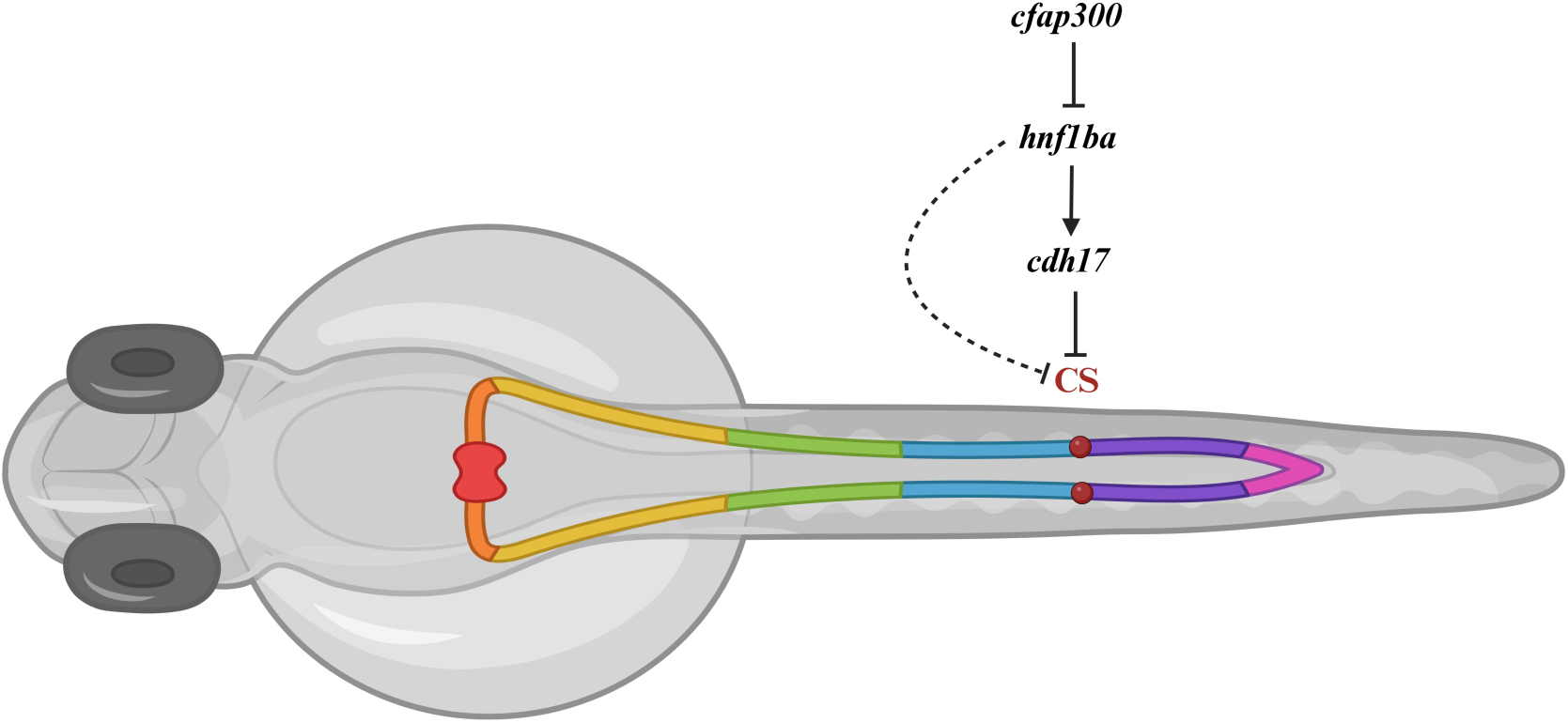
A model depicting the function of *cfap300* during Corpuscle of Stannius (CS) gland formation. The normal function of Cfap300 is to downregulate *hnf1ba*. In the *cfap300* loss-of-function mutant, *hnf1ba* transcription is upregulated, thus upregulating its target *cdh17*, which leads to inhibition of CS gland formation. Hnf1ba may regulate the transcription of genes other than *cdh17*, which may also contribute to CS gland formation.

This study revealed that *cfap300* zebrafish mutants have no major cilia-related phenotypes. In contrast, human CFAP300 mutations lead to PCD. One of the major impacts of PCD is infertility due to reduced sperm motility. The *cfap300* mutants grow to adulthood and produce progeny at par with their WT counterparts, although the sperm of the *cfap300* mutant adults show slow motility with low genetic penetrance (Supplementary Videos 1 and 2). *cfap300* has a strong expression in the pronephros, but the mutants and the morphants have no impact on the pronephros segmentation, MCC formation or pronephros function (Supplementary Figure 4). This indicates that the lack of *cfap300* function in zebrafish mutants may be partly compensated for by other DNAAF, which needs further investigation. It is also tempting to postulate that the role of Cfap300 in the transdifferentiation and CS gland formation may reflect a cilia-independent function, and future work will clarify this. DNAAFs are essentially chaperons that are responsible for assembling protein complexes and it is plausible that Cfap300 may act as a chaperon in signalling networks that control CS development. RA, Notch and FGF signalling pathways are known regulators of pronephros segmentation and contribute to CS gland formation (12, 18, 19). Wnt/Beta-catenin signalling is known to regulate different aspects of pronephros development and patterning (41). The role of Cfap300 in these signalling pathways in the context of CS development and transdifferentiation requires further investigation.

CS gland formation also involves the extrusion of the transdifferentiated cells out of the nephric epithelium. We have observed that the small number of cells that transdifferentiate from the tubule epithelium are not efficiently extruded in the *cfap300* mutants. Cell extrusion is a property of the epithelium that maintains cell homeostasis, where undesired cells are squeezed out from the epithelium without creating a gap, thus preserving the intact barrier property. Being the first line of defence barrier, epithelial cell encounters the highest stress of cell turnover. This turnover stress is an inclusion of a high rate of cell death, overcrowding, replicative stress, pathogen infection, oncogenic mutation, epithelial cell transition or transdifferentiation (42–45). To maintain a constant cell number, epithelium extrudes above mentioned cells by apical cell extrusion (ACE) or basal cell extrusion (BCE) by the coordination of both extruding and neighbouring cells. The mechanisms employed by CS cells for extrusion are not well studied, but are likely to employ some of the known mechanisms of extrusion seen in other biological contexts. At later stages of development (6 dpf and later), a small CS gland can be seen in the *cfap300* mutants (data not shown), further strengthening the idea that transdifferentiation and extrusion processes are tightly regulated processes.

In conclusion, we have uncovered a novel role for a DNAAF in transdifferentiation and organogenesis of an endocrine gland. Our data indicates that Cfap300 may not regulate this process through a classical ciliary function, but through a transcriptional adhesion pathway involving *hnf1ba* and *cdh17*.

## Materials and Methods

### Zebrafish husbandry and ethics statement

Zebrafish were maintained in a circulating system with a 14-hours light and 10-hours dark cycle at 28⁰C ± 0.5. The zebrafish strains, Albino and Tübingen (Tü) were used in all experiments. Embryos were grown in E3 medium at 28.5°C and staged according to the standard protocol (46). All the experiments were approved by the Institutional Animal Ethics Committee (ILS/IAEC-250-AH/FEB-22).

### Total RNA extraction and cDNA synthesis

The total RNA from zebrafish embryos/larvae was extracted using MagSure all RNA isolation kit (RNA Biotech, India) or Direct-zol RNA miniprep kit (Zymo Research, USA) according to the manufactures protocol. Subsequently, equal amount of RNA was taken and the cDNA was synthesised using SuperScript IV First-Strand Synthesis System with oligo (dT)18 (Thermo Fisher Scientific). Finally, the quantitative measurements were taken using in Qubit 4 fluorometer (Qubit™ ssDNA Assay Kit, Thermo Fisher Scientific).

### Probe synthesis for whole-mount *in situ* hybridization (WISH)

The DNA templates required for the probe synthesis (*cfap300*, *cdh17*, *xirp2a*, *gata3*, *fgf23* and *hnf1ba*) were amplified with gene-specific primers (Supplementary Table 1) and Phusion polymerase using cDNA obtained from wild-type (WT) zebrafish embryos. The PCR products were cloned into the PCR-Blunt II-TOPO vector (Thermo Fisher Scientific) and the sequences were verified by Sanger sequencing. The plasmids were linearized and DIG or Fluorescein labelled probes were synthesised using SP6 or T7 RNA polymerases (Supplementary Table 1) (47).

### WISH

The WISH was carried out following the protocol modified from Thisse and Thisse, 2008 (47). In brief, the embryos were fixed in 4% paraformaldehyde (PFA), washed in PBS, transferred to 100% methanol and stored at –20°C a minimum for 2 hours. The embryos were rehydrated in PBST, permeabilised with proteinase-K and refixed in 4% PFA. The embryos were incubated in hybridisation buffer for 4 hours (without the probe), then the probe was added and kept at 65°C for overnight. The next day, the embryos were washed with formamide/SSC buffer at 65°C to remove unbound probes and transferred to MABT (0.1% Tween-20 in 1X MAB) at RT. Then the embryos were transferred to blocking buffer (10% fetal bovine serum (FBS) and 2% blocking reagent (Roche 1109617600) in MABT, 2-3 hours at RT). Then it was incubated overnight with alkaline phosphatase (AP) conjugated anti-digoxygenin (DIG) or anti-fluorescein antibodies at 4°C. On the third day, gene transcripts were detected by developing with BM purple (Roche 11442074001). The reaction was stopped by adding PBS and fixed with 4% PFA kept at 4°C for overnight. To check the expression of two genes in the same embryo, double *in situ* hybridisation was carried out by incubating in both DIG and fluorescein labelled probes targeting two different gene transcripts during the hybridisation step. After detection of the first transcript with BM purple, AP activity was deactivated with methanol, re blocked, and incubated with the second antibody respective to the second probe. The next day, the second RNA probe was visualized using INT-BCIP (Roche 11681460001) and then the reaction was stopped as described above.

### Fluorescent in situ hybridization (FISH)

For two colour FISH, the embryos were hybridized with Fluorescein or DIG labelled riboprobes and incubated with horseradish Peroxidase (POD) conjugated antibodies. The probe was detected using fluorescence substrate from the TSA Plus Fluorescein Kit (NEL741001KT). Then, the first POD antibody was inactivated by the addition of 1% H202 (prepared in methanol). Then the embryos were incubated with POD conjugated antibody against the second probe. The second probe was visualized by adding TSA Plus TMR Solution (NEL763001KT) followed by PBST wash, cleared with 75% glycerol and kept at 4°C for overnight (48).

### Histology

WISH-stained embryos were fixed in 4% PFA for 1-2 hours at room temperature followed by PBS wash and then transferred to 30% sucrose solution (prepared in PBS) for overnight at 4°C. The next day, the embryos were incubated in a 1:1 mixture of OCT medium and 30% sucrose solution for 30 minutes at RT and stored at –80°C. Then 10μm-sized cryo-sections were taken and mounted on a glass slide using a cryotome (Thermo Fisher Scientific).

### TALEN

The TALEN target site for *cfap300* was designed using TALEN-T software selecting two 18 bp sequences in exon 2 as the left (5’ ACAGCACAAGCCTTCAAC 3’) and right (5’ CCAGCCTTATAGAAGCAA 3’) binding sites (https://tale-nt.cac.cornell.edu/) (49). TALEN binding domains were assembled using the Golden Gate cloning method, where RVD repeats 1–10 in pFUS_A and11–17 in pFUS_B1 were cloned using BsaI and T4 ligase. The positive clones were confirmed with universal primers and they were combined with the 18th RVD plasmid (pLR-HD for left, pLR-NI for right) and destination vectors (pCS2+Tal3-DDD and pCS2+Tal3-RRR) using BsmBI (50, 51). Finally, the constructs were confirmed by PCR and subsequently used for *in vitro* mRNA synthesis. The TALEN left arm containing pCS2+-Tal3-DDD and right arm containing pCS2+-Tal3-RRR were linearized with Not I and the mRNA was synthesised using SP6 mMessage mMachine kit.

Both left and right TALENs were co-injected into single-cell zebrafish embryos and the injected embryos were checked for mutation efficiency by high-resolution melt (HRM) analysis and heteroduplex (HD) assay at 48hpf. These fish were grown up to adulthood, and the F0 founders were identified and crossed with wild-type fish to generate F1 heterozygous mutants. The DNA sequencing of F1 heterozygous adults showed the presence of four different kinds of mutations in the *cfap300* gene. These mutations include the deletion of 7, 8, 10 and 26 nucleotides. To generate homozygous mutants, heterozygous fish with deletion of 26 nucleotides from 340-365 of NM_001077345.3 (del26) were crossed with each other, grown to adulthood and homozygous fish were identified by HRM and HD.

### Design and synthesis of guide RNA (gRNA)

The target sites for the gRNA synthesis were identified for the following genes *cdh17*, c-*myc* and *hnf1bb* using CRISPR RGEN tools (http://www.rgenome.net/cas-designer/). For *cdh17*, the targets sites were designed against exon-4 5’GGAGATCCTGACCTTATCTT3’ and exon-8 5’GATTGTTCGGGCTGAGGATT3’. For c-*myc* the targets sites were designed at exon 1-5’ GAGACAGTCGCTCTCCACCG 3’ and exon 2-5’ GGTCATGCCGCGTTGACGGA 3’. For *hnf1bb*, two target sites were chosen at exon 3-5’ GAGCAGCGGTAGGAGATGAG 3’ and exon 9-5’ GAAGATTTCATCCCCTCAGTT 3’. The target sites and the universal oligo were synthesised from IDT. The DNA template was prepared and gRNA synthesized using *in vitro* transcription (52).

### High-resolution melt curve analysis (HRM) and heteroduplex mobility assay (HMA)

The genomic DNA was isolated from the zebrafish embryo/adult fin by incubating the single embryo/adult (52).The target sites were amplified using the site-specific primers (Supplementary Table 1) using 2X SYBR® Green JumpStartTM Taq ReadyMix. During melt curve reaction was performed from 60-95°C with increment of 0.2°C and fluorescence was recorded at every 0.2 sec. Then, PCR amplicons were denatured at 95°C and reannealed at RT. Then, the samples were resolved in 12% polyacrylamide gel at120V for 90 minutes and gel images were taken using ChemiDoc (Biorad).

### Molecular cloning

For the mutation identification in target genes after gene knockout experiments using TALEN and CRISPR, the target sites were amplified and cloned into pGEM®-T Easy Vector Systems. To make zebrafish *cfap300* (818 bp) expression construct, the target region was amplified from cDNA using a forward primer containing NCoI and the reverse primer containing XhoI restriction enzyme sites with an HA tag at the C-terminus. Then it was subjected to restriction digestion and ligated into the pCS2 + vector. After that the transformation was carried out using DH5α cells and cultured in LB media. Finally, the plasmid DNA was isolated by using QIAprep Spin Miniprep Kit (27106) and stored at –20°C.

### *In vitro* transcription

pCS2+-*cfap300* vectors were linearized with Not I and the mRNA was synthesised using SP6 mMessage mMachine kit. pT3TS-nlsCas9nls vector was linearized with XbaI and mRNA was synthesized using T3 mMessage mMachine kit (Invitrogen). For the gRNA synthesis, annealed gRNA oligos were used as templates and gRNA was synthesised using Invitrogen^TM^ MEGAScript^TM^ T7/SP6 kit and purified using the phenol and chloroform extraction method.

### Morpholino design

The antisense morpholino oligonucleotide was designed and purchased from Gene Tools. *cfap300*-ATG morpholino (*cfap300*-MO, 5′ TGCCTATTGTTGTTGTTACCATGAC-3′) was designed to block translation of *cfap300* mRNA and a mismatch morpholino (*cfap300*-MM) 5′ TGCaTATTaTTaTTaTTACCATaAC-3′ with five base mismatches (small letters) was used as a negative control. For *hnf1ba* a previously reported morpholino (5’ CTAGAGAGGGAAATGCGGTATTGTG 3’) was used in our study (53). Morpholinos were dissolved in nuclease-free water to make a 1 mM stock, diluted to 0.25mM and 1nl was injected into each embryo at the 1-2 cell stage.

### Microinjection

Microinjection was performed at single cell stage of zebrafish embryos for all following experiments using a Femtojet microinjector (Eppendorf). 100 pg/nl of right and left arm mRNA was used for the microinjection for the TALEN-mediated mutagenesis. The gene knockout by CRISPR mutagenesis was achieved by injecting 1nl of gRNA and nCAS9n mRNA mixture in the concentrations of 25 ng/μl, 100 ng/μl, respectively. *cfap300* mRNA was injected at 150 pg per embryo for rescue experiments.

### Quantitative Reverse Transcription PCR (RT-qPCR)

Comparative gene expression analysis of *cdh1*, *cdh6*, *cdh17* and *cfap300* and transcription factors *cjun*, *ppargc1a*, *c-myc, hnf1ba* and *hnf1bb* were carried out using RT-qPCR. The experiment was performed in QuantStudio™ 5 Real-Time PCR System (Tm 60°C, 40 cycles) using GoTaq® qPCR Master Mix (Promega). Relative gene expression levels were calculated using the ΔΔCt method, normalising to housekeeping gene *ef1α*.

### Digital PCR (D-PCR)

The expression levels of *fgf23*, *cdh17* and *hnf1ba* mRNA were quantified by QIAcuity One D-PCR system. The reaction mixture was prepared by adding cDNA, target specific primers and EvaGreen master mix from the QIAcuity^TM^ EG PCR Kit (Qiagen). The PCR reaction was carried out with the following cycles: Initial denaturation 95⁰C-2 min, then 40 cycles of denaturation at 95⁰C for15 seconds, annealing at 62⁰C for 30 seconds and extension at 72⁰C for 25 seconds and final extension at 40⁰C for 5 min. Images were acquired at an exposure of 300 milliseconds and a gain of 6 in the green channel. The result was analysed using QIAcuity Software Suite (Qiagen). Relative gene expression levels were normalized to the housekeeping gene *ef1α* and *b2m*.

### Transcription Factor Identification

To identify the transcription factors binding to the *cdh17* promoter, an in-silico promoter analysis was carried out using ALGGEN PROMO software (http://www.lsi.upc.es/~alggen). After the analysis it was found that 234 transcription factors can bind to *cdh17* promoter. Spatial expression of these TFs during zebrafish development was checked at the Zfin.org database to identify the TFs that are expressed in a spatially overlapping manner with the *cdh17* in the zebrafish pronephros. The TFs *cjun*, *ppargc1a*, *hnf1ba*, *hnf1bb* and *c-myc* were found to be expressed in a complete or partially overlapped fashion with *cdh17* and were taken for further analysis.

### Microscopy

The zebrafish embryos were mounted in 75% glycerol and the bright-field and fluorescence images were taken using Leica M205 FA stereo microscope with identical magnification and exposure settings for the all the samples.

### Pronephros function assay

For pronephros function assessment, 70 KDa dextran was injected into the common cardinal vein of 54 hpf zebrafish embryos. Accumulation of dextran was measured by measuring the fluorescent intensity at the heart, using ImageJ software at 0, 5 and 24 hours post-injection. Then the percentage of fluorescent intensity was calculated to measure the clearance by the pronephros(54).

### Sperm motility assay

Sperm collection and motility test were carried out following Kikkawa and Yamaguchi, 2019 (bio-protocol.org/prep21). Spermatozoa were collected in ice-cold Hank’s buffer, then diluted in 0.2X Hank’s buffer and used for video microscopy. Sperm motilities were observed under bright-field conditions using Celldiscoverer 7 (Zeiss). The video was obtained at 20X magnification and played at 12fps.

### Statistical analysis

The statistical analysis of relative gene expression was performed using an unpaired t-test and a 2-way ANOVA method in GraphPad Prism 8.0.1. and the data were presented as mean ± standard deviation (SD). The graphs representing the statistical significance p<0.05 (*), p<0.01 (**) and p<0.001(***) or as mentioned in the figure legends. For rescue experiments, the significance was measured by implementing the Cochran–Mantel–Haenszel (CMH) test. The analysis was performed using IBM/SPSS software and a p<0.001 was considered significant.

## Supporting information

Movies of free-swimming spermatozoa of WT

Movies of free-swimming spermatozoa of cfap300 mutant

## Acknowledgements

We thank Suryasikha Mohanty for her help with the experiments, Satyajit Behera and Subhangi Mohanty for maintaining the zebrafish facility. We thank Dr Chinmoy Patra, ARI, Pune for discussions and critical reading of the manuscript and Dr Vinoth S. for help with manuscript preparation. This work was partially supported by a SERB-EMR grant (EMR/2016/003/780) and a DBT grant (BT/PR45460/MED/12/952/2022) to RKS, and intramural funds from ILS, which is an institute of BRIC, DBT, Government of India. UN is a recipient of the DST-Inspire fellowship (IF180156) and KS is a UGC-SRF (221610148651). Figure 9 was created with BioRender (Agreement number-BA28TETTCD).

## Author Contributions

RKS conceptualised the project. PB and UN generated the mutants and reagents. WISH and fluorescent WISH were carried out by UN, KS and PB. UN and KS performed RT-qPCR and RT-digital PCR experiments. UN, KS and PB analysed the data and all authors prepared the manuscript.

## Figure Legends

**Supplementary Figure 1:**
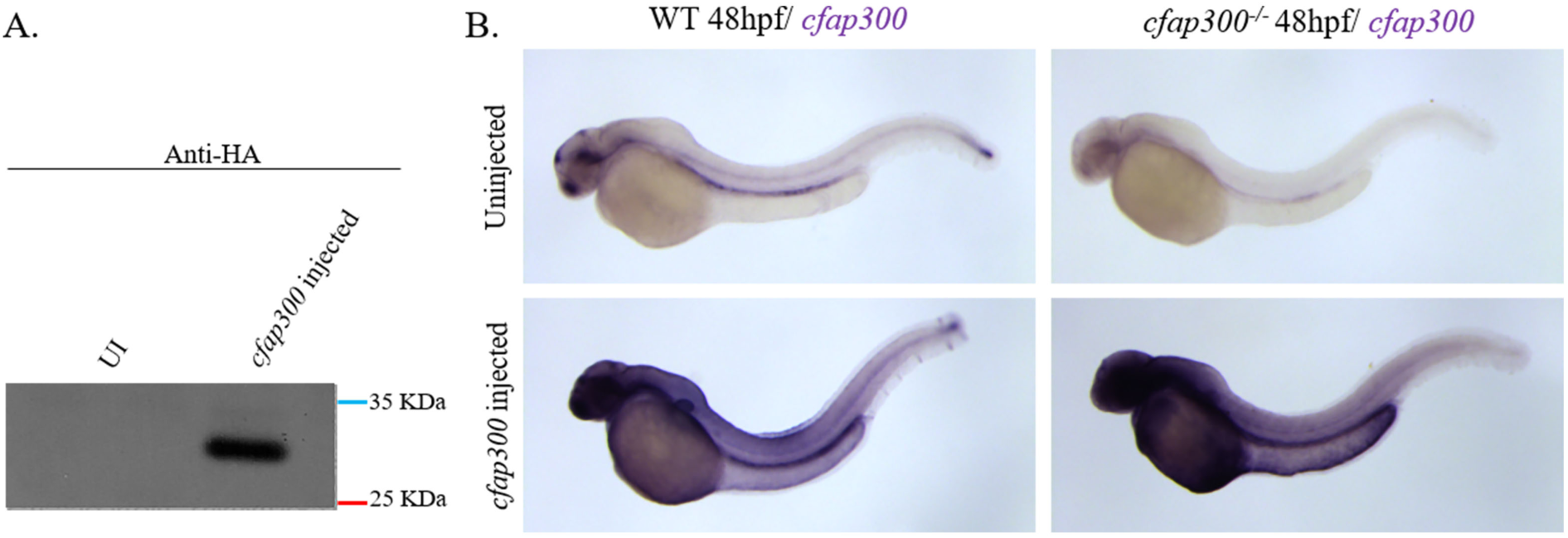
Expression of the *cfap300*-*HA* mRNA. **(A)** Western blotting showing the expression of Cfap300-HA. **(B)** Microinjection of *cfap300-HA* mRNA in WT and *cfap300* mutants shows high and ubiquitous expression.

**Supplementary Figure 2:**
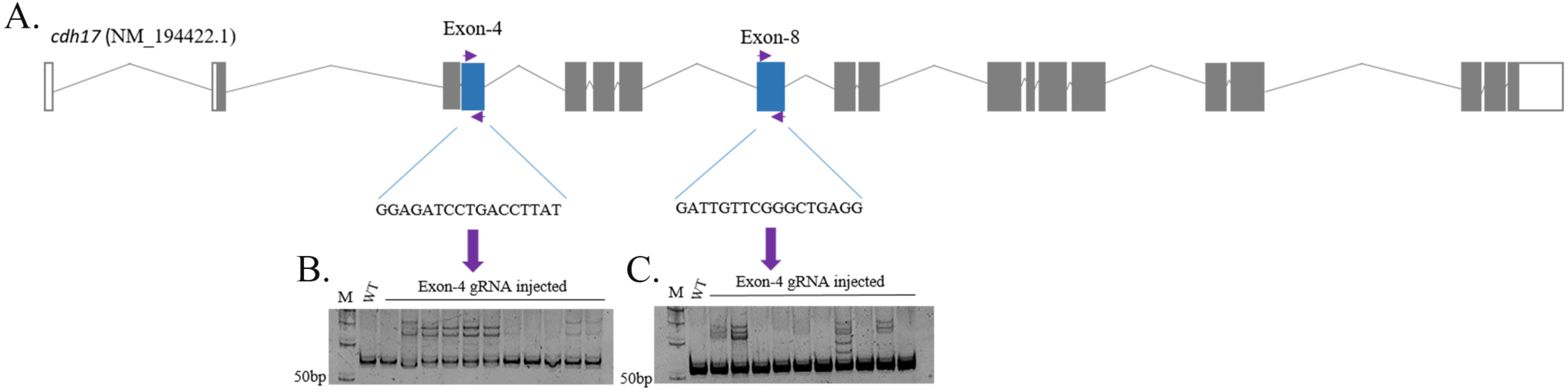
Efficiency of *cdh17* gRNA. **(A)** gRNA was designed against exon-4 and –8, and these gRNA and Cas9 mRNA were microinjected into one-cell stage zebrafish embryos. **(B – C)** Both gRNAs can induce high efficiency mutagenesis as seen by HD.

**Supplementary Figure 3:**
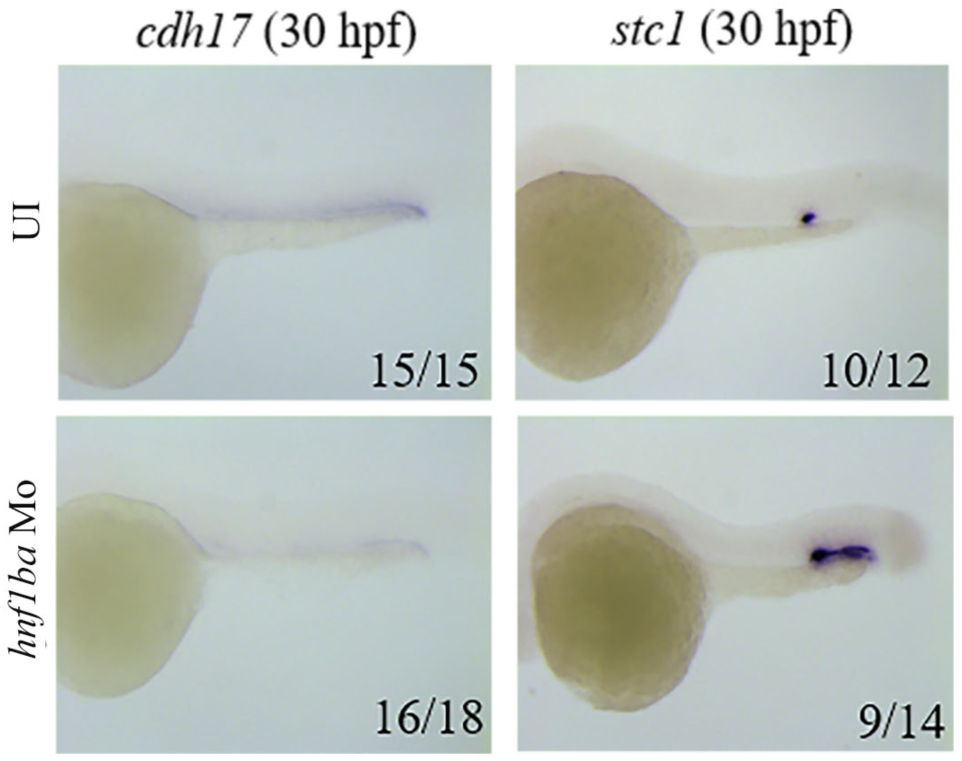
Effect of *hnf1ba* knockdown on *cdh17* and *stc1* expression. WISH shows knockdown of *hnf1ba* reduces *cdh17* expression and enhances *stc1* expression.

**Supplementary Figure 4:**
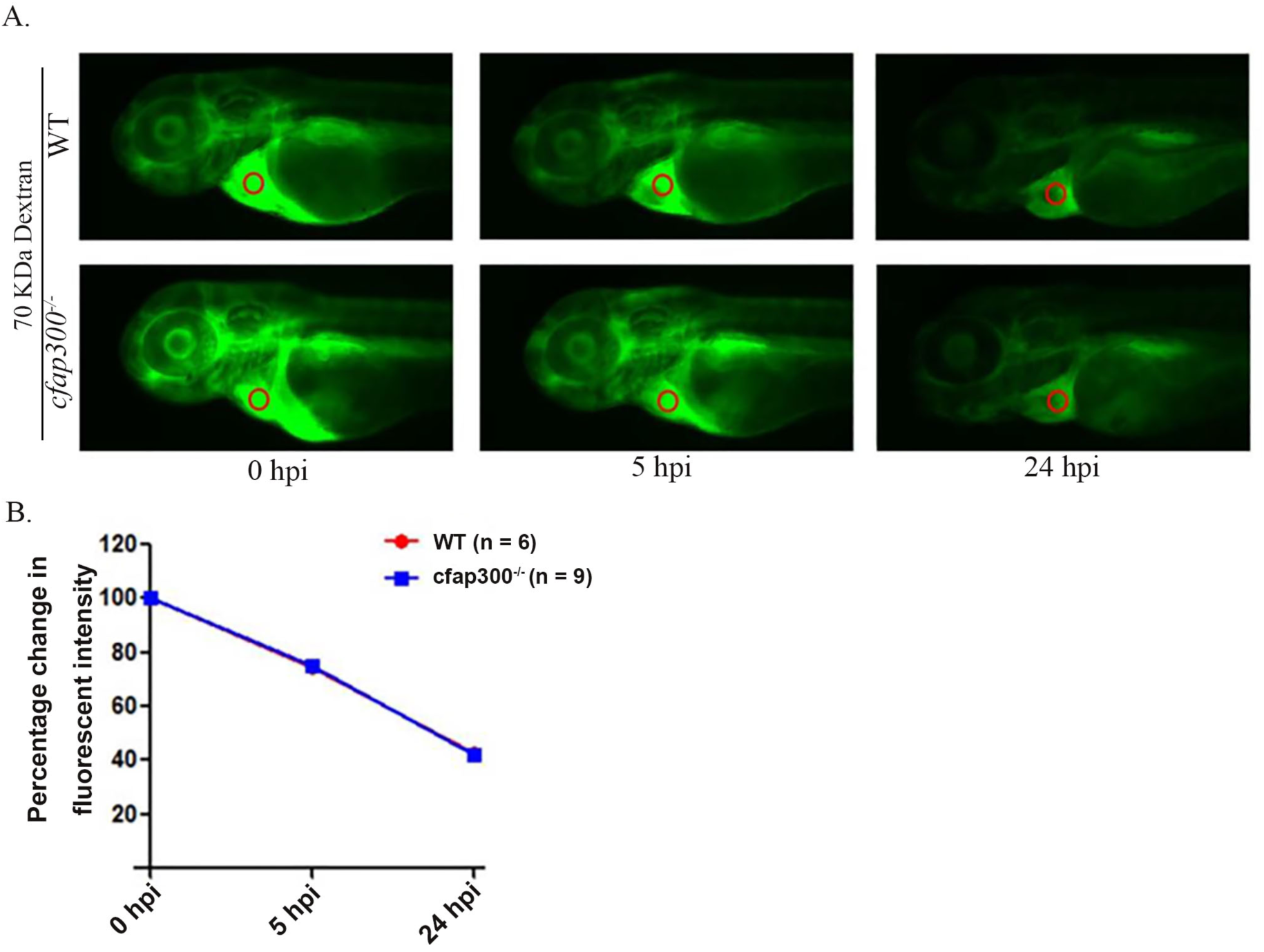
Pronephros function is not affected in *cfap300*^−/−^ embryos. **(A-B)** Clearance of 70 kDa dextran injected into the common cardinal vein of 54 hpf WT and *cfap300^−/−^* embryos. **(A)** Representative picture of dextran-injected WT and *cfap300^−/−^* embryos. The fluorescence intensity of a defined area (Red circle) was measured in each embryo at 0, 5 and 24 hours post-injection (hpi) **(B).** The percentage change in fluorescent intensity was plotted on a graph.

**Supplementary Video 1 and 2:** Movies of free-swimming spermatozoa of WT and *cfap300* mutant, filmed by a high-speed camera and played at 12 fps. Scale bar: 50 μm

**Supplementary Table:**
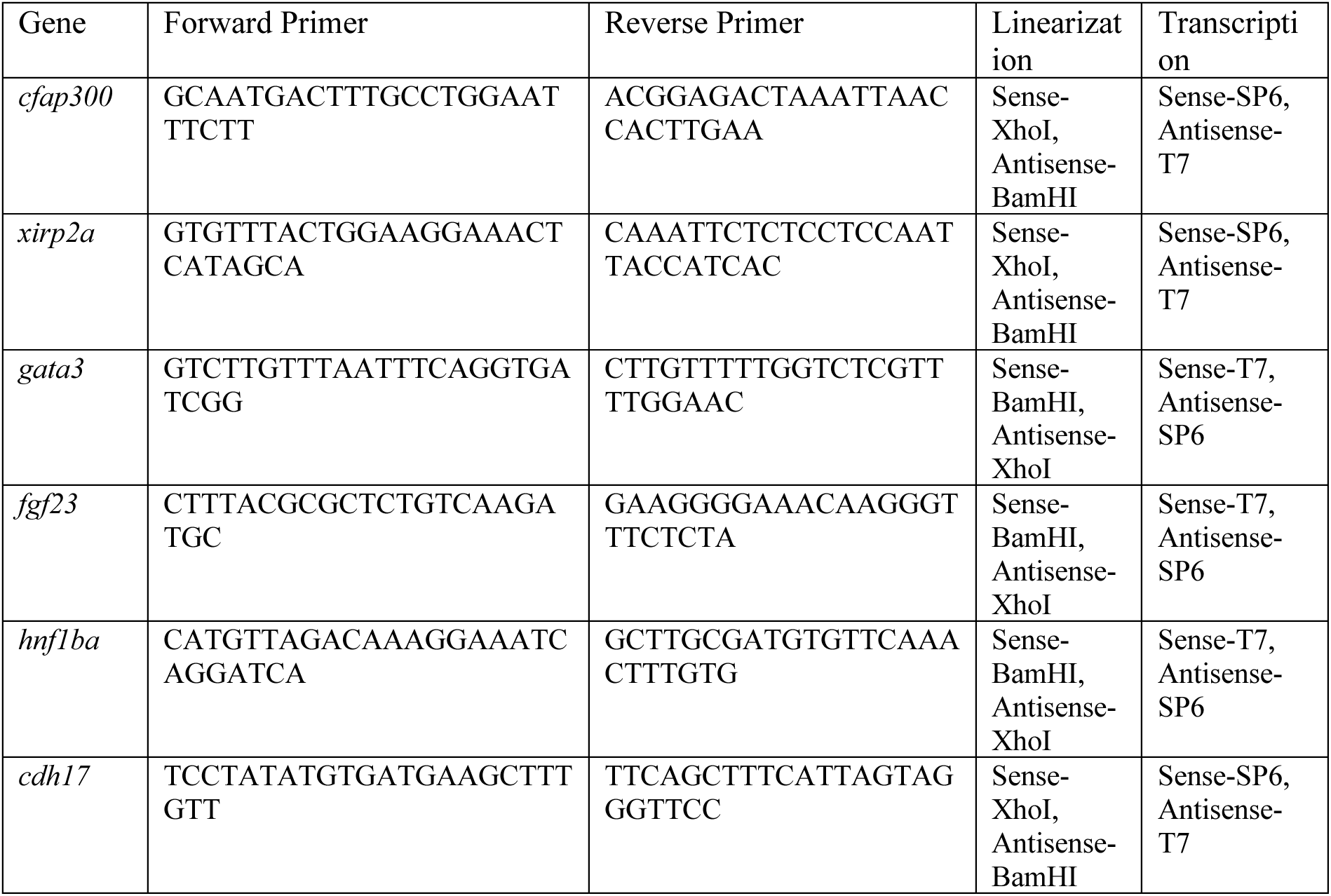
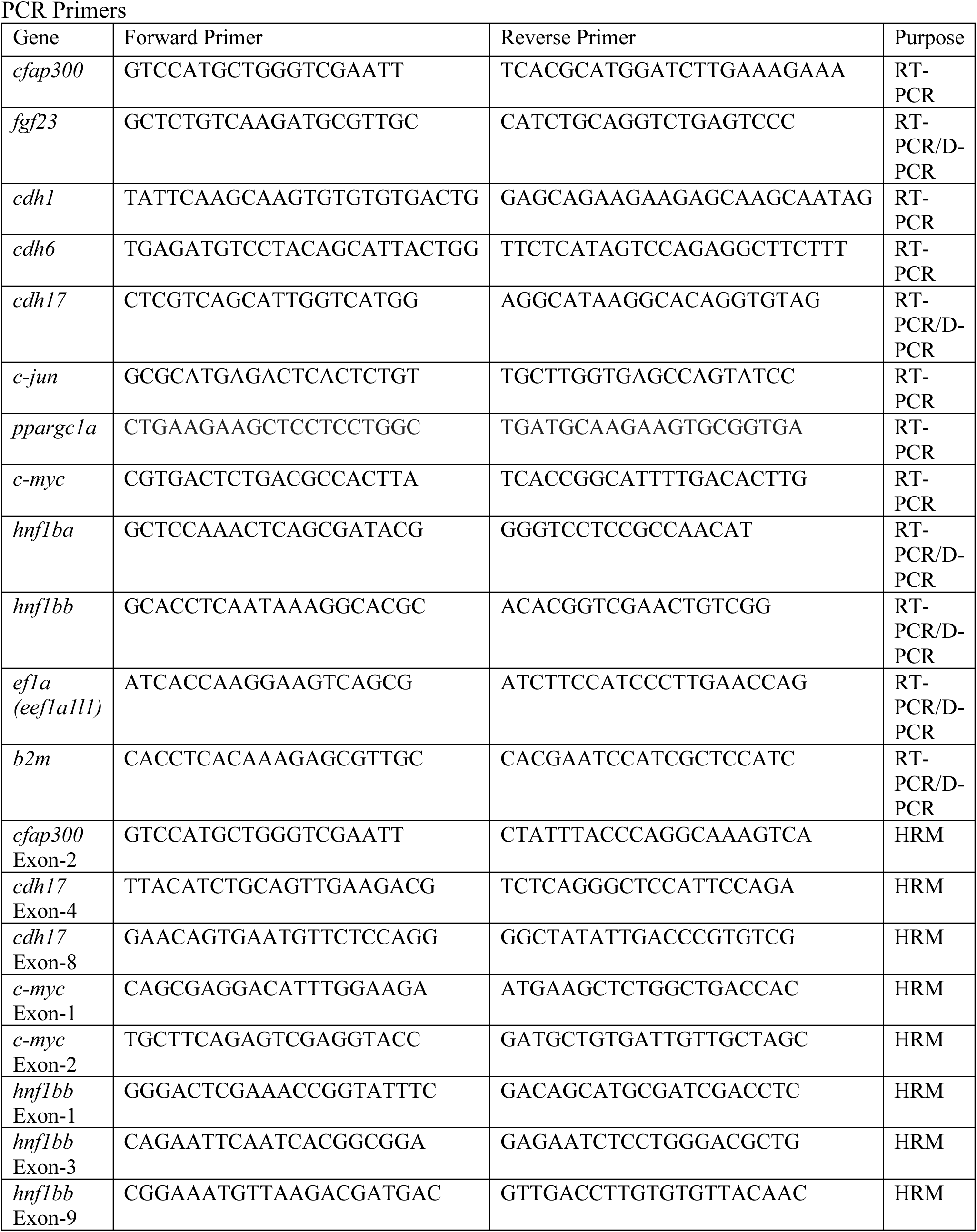
Oligonucleotides used in PCR. Primers used to clone WISH probes

